# A Proof Of Concept For A Syndromic Surveillance System Based On Routine Ambulance Records In The South-west Of England, For The Influenza Season 2016/17

**DOI:** 10.1101/462341

**Authors:** Thilo Reich, Marcin Budka

## Abstract

The introduction of electronic patient records in the ambulance service provides new opportunities to monitor the population. Most patients presenting to British ambulance services are discharged at scene. Ambulance records are therefore an ideal data source for syndromic early event detection systems to monitor infectious disease in the prehospital population. It has been previously found that tympanic temperature records can be used to detect influenza outbreaks in emergency departments. This study investigated whether routine tympanic temperature readings collected by ambulance crews can be used to detect seasonal influenza. Here we show that these temperature readings do allow the detection of seasonal influenza before methods applied to conventional data sources. The counts of pyretic patients were used to calculate a sliding case ratio (CR) as a measurement to detect seasonal influenza outbreaks. This method does not rely on conventional thresholds and can be adapted to the data. The data collected correlated with seasonal influenza. The 2016/17 outbreak was detected with high specificity and sensitivity, up to 9 weeks before other surveillance programs. An unanticipated outbreak of E. coli was detected in the same dataset. Our results show that ambulance records can be a useful data source for biosurveillance systems. Two outbreaks caused by different infectious agents have been successfully detected. The routine ambulance records allowed to use tympanic temperature readings that can be used as surveillance tool for febrile diseases. Therefore, this method is a valuable addition to the current surveillance tools.

## 1 Introduction

An increasing amount of digital data is being generated every day in all domains of human activity. The health services are not an exception and have also experienced an expansion of their digital footprints. One recently digitised area of the health service are patient records collected by ambulance crews. These have previously been handwritten and therefore difficult to access. This digitisation has created a large prehospital data source that to date is mostly untapped.

The South-Western Ambulance Service NHS Foundation Trust (SWASFT) introduced electronic Patient Care Records (ePCR) in March 2015 using a staggered approach. The ePCR allows to access and monitor all data recorded near real-time. SWASFT has a high non-conveyance rate of 57.45% (2013-2014), which is the highest in the UK [1]. This allows to monitor the community rather than the hospital population.

Most infections result in pyrexia [2,3], therefore body temperature-based surveillance systems are unspecific but sensitive to virtually any pyrexia causing disease.

In the past, temperature screening has been applied during outbreaks of infectious diseases such as the Severe Acute Respiratory Syndrome (SARS) [4–6] and was routinely used to identify carriers at airports. It has also been demonstrated that the monitoring of body temperature on its own, allowed to detect an outbreak of seasonal flu in an emergency department in Boston (US) [7].

Such a syndromic surveillance system should ideally be deployed within the community before patients are admitted to hospitals. Therefore, the ambulance service is ideally placed to provide such a detection system [8].

Most surveillance systems rely on user set thresholds to detect outbreaks [9]. This study will demonstrate that it is possible to use patient records collected by ambulance crews to detect disease outbreaks, solely based on tympanic temperature readings, without relying on user-set thresholds. The key objectives of this study are:

- To establish if the prehospital tympanic temperature readings mirror the seasonal influenza peak during the 2016/2017 season.
- The evaluation of a method adapted from Singh et al. (2010) [10] using Case Ratios (CR) and its applicability as Early Event Detection (EED) system when applied to prehospital tympanic temperature readings.

## 2 Methods

### 2.1 Data Extraction

The retrospective ePCR dataset was provided by SWASFT Clinical Information and Records Office and was extracted from a database using queries for the counties of Cornwall and Devon. Records without valid postcodes were excluded (1.8% of 375,740). The variables used are the record creation date, tympanic temperature and age.

All data were collected during routine patient contact of ambulance crews. As the nature of this study was a clinical audit and no identifiable patient data was extracted, an ethical approval was not required [11]. The aim of this work was to evaluate whether these data can be used for a surveillance system in the future. As the data is stored digitally it could be accessed near real time which would be a requirement of such as system.

The most commonly used temperature probes within SWASFT are the Braun ThermoScan 7 IRT6520 and ThermoScan 5 IRT4520. Both devices have a measurement range of 34-42.2° C with an accuracy of 0.2° C between 35.5-42° C and 0.3° C outside this range [12,13]. As the data included temperature readings outside the range of the device capabilities, records indicating physiological hypothermia (32° C) [14,15] and hyperpyrexia (42° C) [7] were excluded.

The most commonly used temperature probes within SWASFT are the Braun ThermoScan 7 IRT6520 and ThermoScan 5 IRT4520. Both devices have a measurement range of 34-42.2° C with an accuracy of 0.2° C between 35.5-42° C and 0.3° C outside this range [12,13]. As the data included temperature readings outside the range of the device capabilities, records indicating physiological hypothermia (32° C) [14,15] and hyperpyrexia (42° C) [7] were excluded.

### 2.2 Data Processing

All data were processed using MATLAB R2017a (The MathWorks, MA, USA).

The human body temperature varies during the day due to changes in metabolism rates. The lowest oral measurement (37.2° C) was found by Mackowiak *et al.* (1992) [16] to be at 6am and the highest (37.7° C) at 4pm. A cohort study conducted by Obermeyer *et al.* (2017) [17] in 35,488 healthy patients confirmed the daily variation and found a normal range of 35.3-37.7° C. This study also established a negligible discrepancy of −0.03° C of tympanic versus oral temperature [17]. Therefore, temperatures between >37.7-42° C, were considered as pyrexia where the upper cut-off accounts for hyperpyrexia [7].

The analysis was performed on records of patients with tympanic temperature readings within the range of 32-42° C. This excluded 0.0306% of the records (i.e. moderate and severe hypothermia as well as hyperpyrexia) – Figure 1B [14].

**Figure 1:**
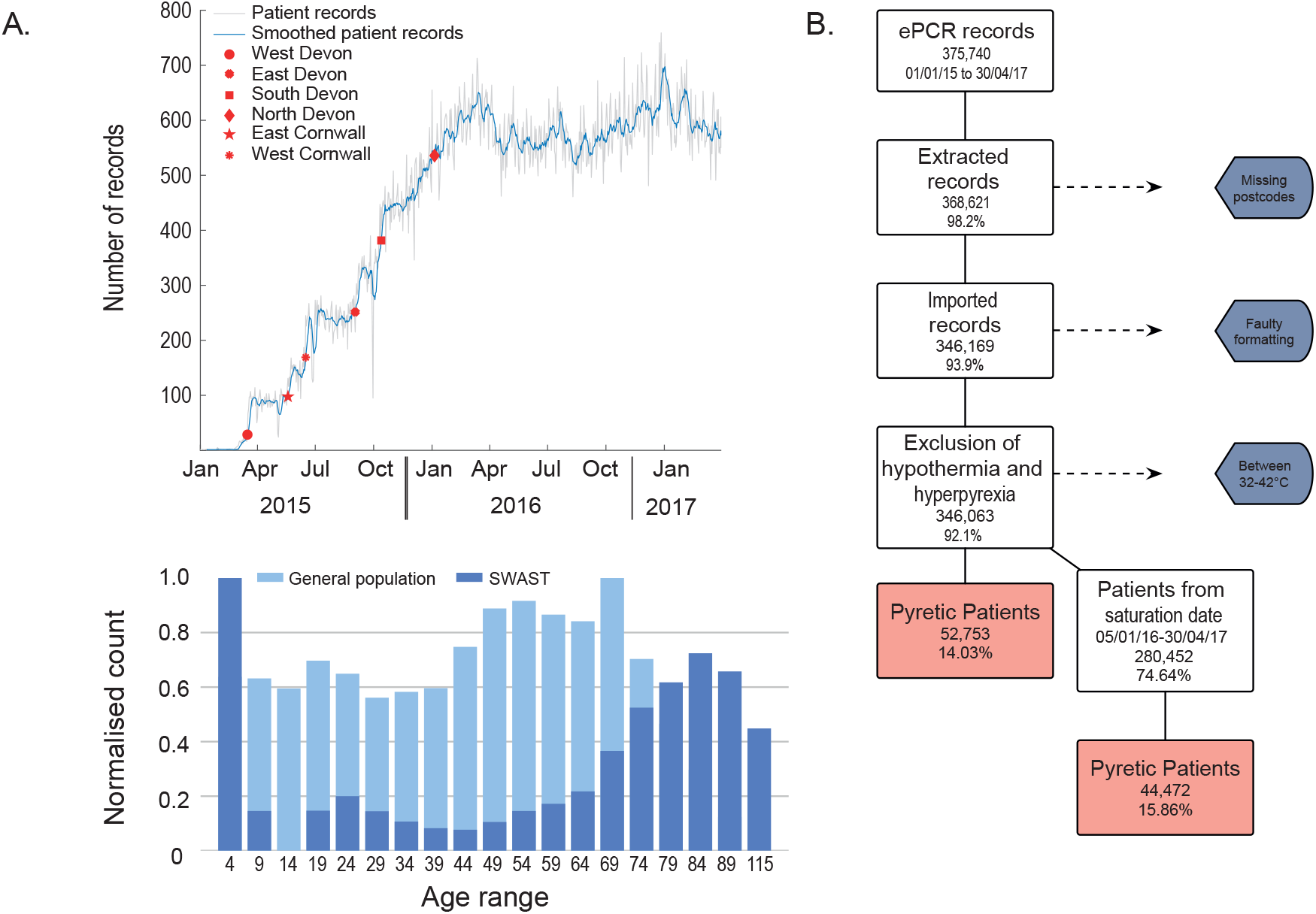
A. Total number of electronic patient records (ePCR) since its introduction within Devon and Cornwall. B. Data loss following exclusion criteria. C. Age distribution for both the general population in Devon and Cornwall in comparison to the patients attended by ambulance crews.

Daily and weekly counts of call volumes and pyretic patient numbers were used as a basis for all following data analyses. Due to the staggered deployment of the electronic devices within SWASFT, the point of data saturation at which all areas were fully operational, was reached on the 05/01/2016 (Figure 1A). Thus, the timeframe was limited to the period between 05/01/2016 and 30/04/2017. As the saturation date, did not capture the start of the 2015/2016 flu season, all detections were run against the 2016/17 flu peak.

### 2.3 Data Smoothing

As the daily patient count varies considerably from one day to another resulting in a noisy time series, the data was smoothed using an Exponential Moving Average (EMA) with an averaging window of 21 days. This window size was chosen because the incubation period of influenza can be up to 3.6 days [18] followed by an onset of symptoms and transmission period of the virus, which can last up to 10 days in hospital [19–21]. This means a patient could be contagious for up to 14 days following infection. Accounting for the incubation period, a secondary patient could show symptoms ~18 days after the infection of the index case. Therefore, the averaging window size was chosen to be 21 days as this allows for some leeway.

An EMA method was used because it weighs data points higher if these are closer to the present compared to samples from the more distant past [9]. This puts emphasis on data from new patients rather than on older data points.

The weekly summed data were used without smoothing as well as an EMA of 3 weeks equivalent to 3 sample points.

### 2.4 Baseline calculation

The weekly sums and the smoothed daily and weekly counts of pyretic patients were binned with bin-counts calculated using the Freedman-Diaconis Rule [22]. The centre of the most frequent bin range was determined and will be referred to as baseline.

### 2.5 Normalisation

Figures showing variables of different scales were normalised using feature scaling to represent the values on a scale between 0 and 1. This is indicated in the figure legend.

### 2.6 Reference data sets

To establish whether the seasonal influenza outbreak is detectable in the ePCR data, these were compared to a reference dataset of weekly Influenza cases in England obtained from the European Centre of Disease Control (ECDC)

1

### 2.7 Calculation of the modified Case Ratio CR_d_

The ability of an infectious agent to spread within a population can be described using the basic reproduction number or R_0_. This value indicates the mean secondary infections caused by each infected host, in a naïve population without immunity against the infectious agent. R_0_ is calculated retrospectively using information about the number of contacts of each infected individual and the resulting secondary infections [23].

Methods exist to estimate R_0_ from the progress of a disease outbreak, which do rely on knowledge about the transmission characteristics of the infectious agent gathered from previous outbreaks [24–27]. As this study aims to demonstrate a method based on temperature readings only caused by any infection this information is not available which is similar to the situation during an unidentified outbreak. Singh et al. (2010) [10] demonstrated that weekly case ratios (CR) can be used as an indirect measure of R_0_ and allow to detect pandemic influenza outbreaks.

The method was adapted in this study by using several different time frames compared to a solely weekly CR, furthermore it was applied to pyrexia numbers rather than specific influenza cases and thus will naturally include infections caused by other infectious agents. To allow to distinguish between different time frames used to calculate the modified CR in this study it is referred to as CR_d_ where *d* represents the chosen time step between observations in days. This means that the time step *d* must be chosen by the investigator to estimate CR_d_ as follows:

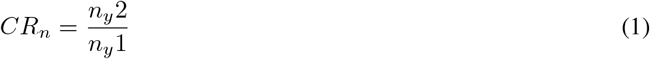

where n_y_ represents the number of pyretic patients at the days of observation with the previously defined time step between observations in days. Thus, n_y1_ represents the first observation and n_y2_ the latest.

Here this method is applied to pyrexia cases as an unspecific substitute for infection, which.

### 2.8 The different mean-CR_d_ depending on window sizes

To establish the effect of different choices of *d*, the ascending area of pyrexia cases peak in 2016/17 was used to calculate a sliding CR_d_ with varying *d* for the ascending slope where pyrexia cases increased (Figure 2).

**Figure 2:**
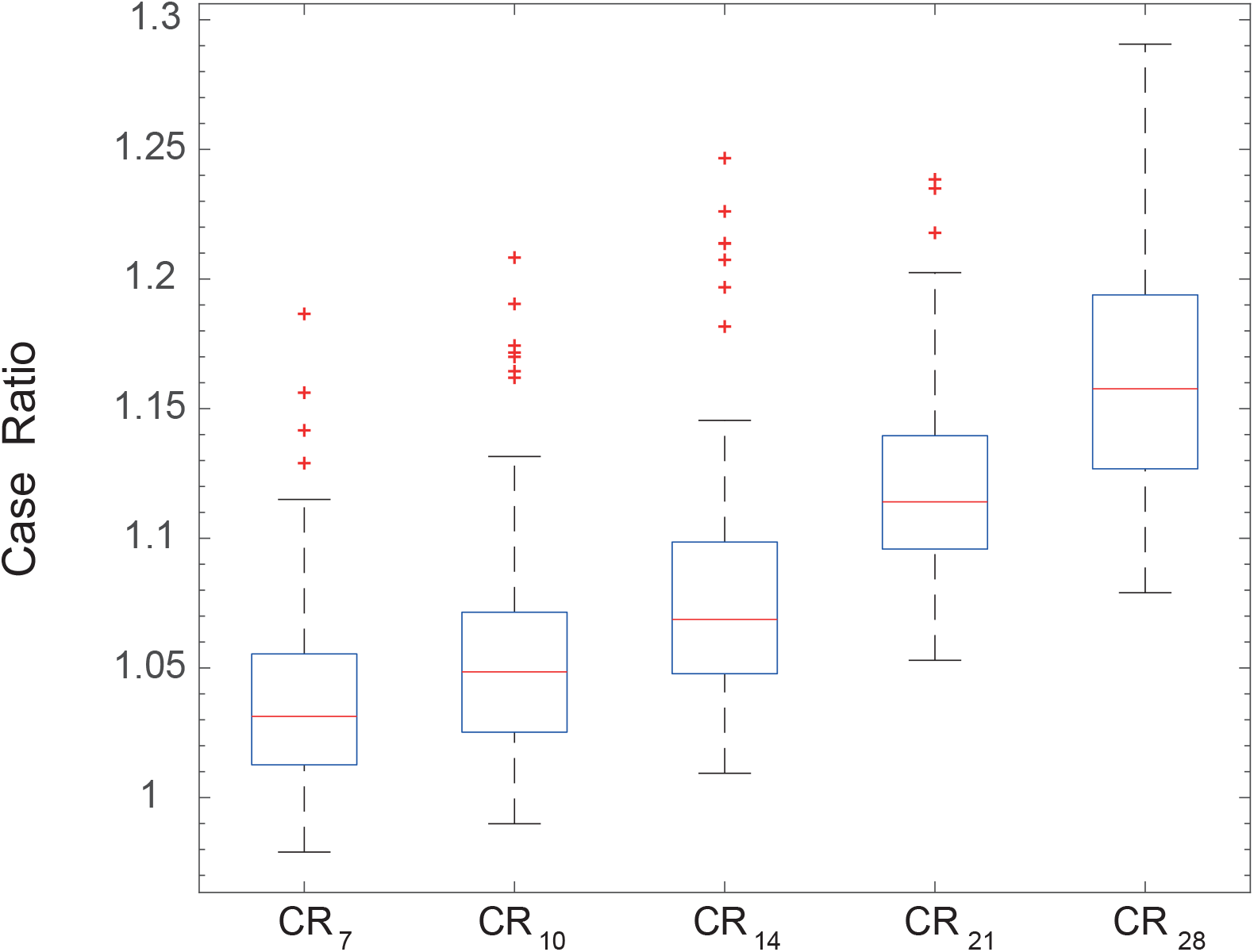
Comparison of different Case Ratios (CR) calculated with different time steps (d).

CR_21_ was chosen here as its values for the ascending slope were greater than 1, thus fitting the assumption that the slope represents an increase in case numbers. Although it included more outliers than CR_14_, it caused less delay than CR_28_, so is effectively a trade-off. To make the approach comparable between daily and weekly counts the same window is used for both sampling rates meaning a window of 21 days or 3 weeks for the daily and weekly counts respectively. The data was tested for auto correlation (not shown) and was found to be auto correlated over the time frame of a year, which would be expected for a seasonal infection. The auto correlation was minor for the 21 day window size and was therefore not further addressed.

### 2.9 Outbreak definition

In this study the outbreak definition is focused on the ascending slope representing an increase in pyrexia case numbers. Therefore, the definition of an outbreak is the persistent transition from CR_d_ <1 to CR_d_ >1 indicating an increase in cases, which remains >1 for at least one week. It has to be noted that outbreak here refers to an increase of pyrexia cases caused by an unspecified infectious agent and could be caused by several agents circulating at the same time.

The delay of one week was chosen because it includes the incubation time, meaning that secondary patients exposed to influenza should have developed pyrexia within one week [18].

Using the CR_d_ as an indication of an outbreak start is based on the assumption that a disease becomes uncontrolled once CR becomes larger than 1 [28]. The final change of CR_d_ to >1, that is not followed by a recovery to <1 within one week until the curve reaches its peak is considered the ascending period of the outbreak.

### 2.10 Estimation of sensitivity and specificity

Both values were calculated from the point at which the CR returned to <1 after the 2015/16 seasonal peak until the peak of 2016/17.

This estimate assumes that the peak occurring simultaneously with the influenza cases detected by the ECDC was a verified event and the unknown peak during summer discussed below was the second verified event see section below. A false positive thus was an increase of the CR followed by a decline to normal levels without a peak in the data provided by the ECDC. If a peak reported by the ECDC fell inbetween an increase of the CR and the subsequent decrease these peaks were deemed to origin from the same event.

## 3 Results

### 3.1 Data characteristics

The dataset from the date of saturation was made up of 54.01% female and 45.99% male patients. The age range was between 0-115 years (one outlier of 864 years was excluded) with a mean age of 59.98 and a median age of 68, where the discrepancy between the two can be explained by a peak in ambulance attendances to infants.

The estimated population in 2016 was 779,834 for Devon and 553,687 for Cornwall. The combined population was 1,333,521 [29]. In comparison the to the estimated age distribution of Devon and Cornwall our data is different as it is skewed towards the elderly and infants as shown in figure 1. This reflects that older individuals and young parents are more likely to require assistance by the ambulance service.

Temperatures recorded were in range of 21-47° C, with a mean temperature of 36.89° C. As this temperature range is unphysiological it was concluded that some data points were caused by input errors. The temperature-based exclusion removed 8 patients (0.0023%) with temperatures of >42° C, 3 of those within the 42-42.2° C range. The lower temperature cut-off removed 98 patients (0.028%) with temperatures <32° C.

To establish whether seasonal influenza was detectable in the dataset, weekly case numbers were compared with weekly sentinel influenza cases recorded by the ECDC in England (see Figure 3B). Sentinel surveillance data is based on a network of selected health care facilities which select patients with symptoms suggesting influenza for laboratory confirmation. The non-sentinel surveillance is a passive system using patient samples for laboratory confirmation of a variety of sources which are not necessarily from patients showing symptoms of an influenza infection [30].

**Figure 3:**
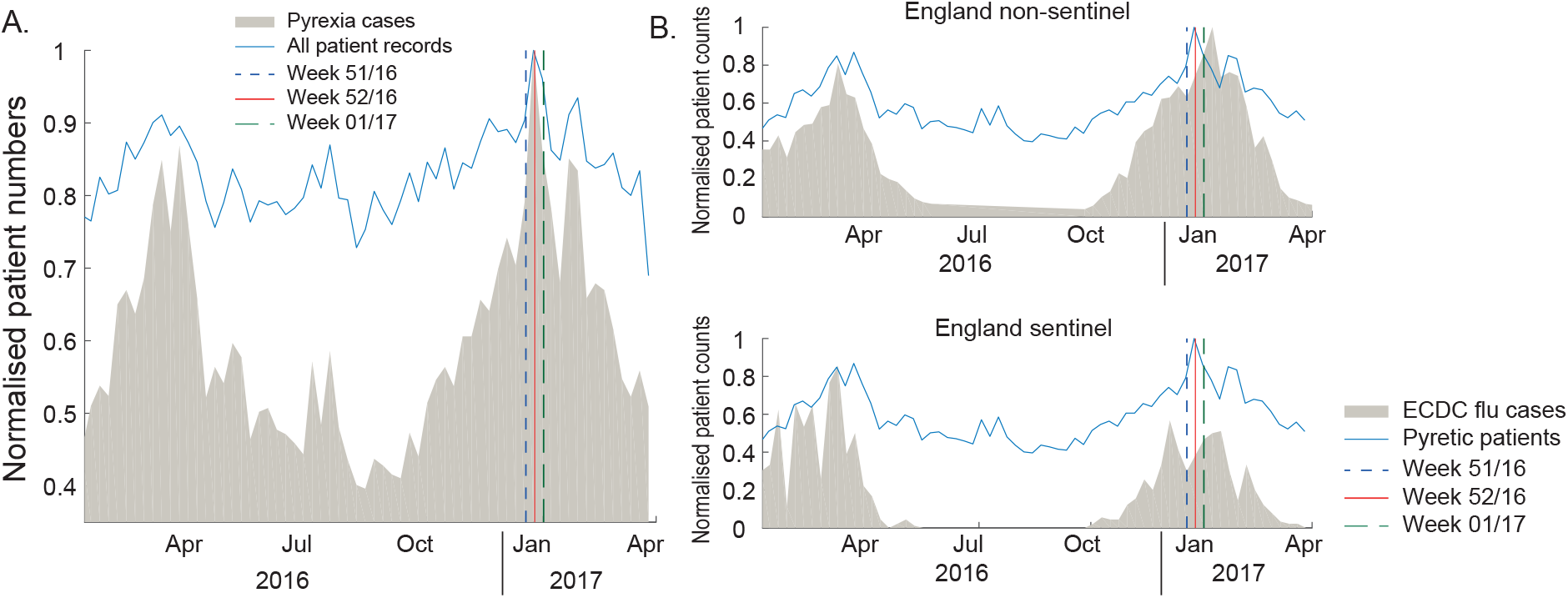
A. Comparison of normalised pyrexia cases and all patient records. B. normalised weekly pyrexia recorded in electronic patient records compared to flu cases in England as reported by the European Centre for Disease Prevention and Control. The vertical lines indicate the detection of the peaks Public Health England for the following syndromes: influenza like illness in week 01/17, Acute Respiratory Illness in week 52/2016.

Both peaks seen for the influenza season 2015/16 and 2016/17 correspond to the data collected by the ECDC. The comparison to the non-sentinel data shows an earlier peak of the ePCR data in the 2016/17 flu season.

This could be explained by the nature of the non-sentinel data, which includes routine patient samples, because the screening is non-targeted.

The ePCR data is mirrored in both the sentinel and non-sentinel laboratory confirmed influenza cases. The ePCR data has a stronger similarity to the non-sentinel surveillance systems which could be explained because all patients are included regardless of symptoms which is also true for the non-sentinel surveillance system. It has to be noted that as the data sources are from different populations and based on different diagnoses. This complicates the proof of causality between the two curves. However, a conducted Granger causality test was significant suggesting that in the absence of a more likely explanation the peak seen in the ePCR data is most likely caused by influenza cases. Some unknown proportion of pyrexia cases will be caused by other infections but if can be expected that the majority was caused by the circulating seasonal influenza virus.

The temporal similarity of our data and the ECDC data shows that the use of temperature as sole indicator of infection allows to monitor unspecific infectious diseases within the community.

A spike in call volumes was seen simultaneously with the peak in pyretic patients and represented an increase of calls by 26.5% compared to the baseline (Figure 3A). This is interesting and could have implications for the preparedness of the ambulance service if such an increase could be predicted.

Public Health England (PHE) monitors influenza cases with different surveillance programs. The data is based on diagnoses from hospitals as well as from GPs. It is separated into Influenza Like Illnesses (ILI) and Acute Respiratory Infections (ARI) [31].

PHE recorded a peak of ILI consultations in week 01/2017, ARI consultations peaked in week 52/2016. This correlates temporally with the peaks seen in the daily summed data (week 01/2017) and the weekly summed data in (week 51/2016), indicating that the seasonal influenza outbreak progresses similarly in both data sets and therefore allowing a direct comparison.

### 3.2 Daily detection

The smoothed daily pyretic case numbers peaked on the 02/01/17 (week 1/2017). The peak was reached with 133 (18.47%) patients of 721 calls (fractions are caused by the smoothing process using the EMA).

The start of the increase in pyrexia cases defined by a final change of the CR_21_ to >1, was detected on the 24/09/16 (week 38/2016) with 70 pyretic patients (11.2% of 633 calls, Figure 4). This value is 6.8% below the baseline (76.2) and within the standard deviation (23.48), which would not be detectable using a threshold method. This start of the seasonal increase of infections was detected earlier than influenza cases by the ECDC, which identified the start in week 46/2016 [32]. However, this might be due to the fact that the sentinel surveillance runs between October and March and thus had not started when our system detected an increase in cases [33]

**Figure 4:**
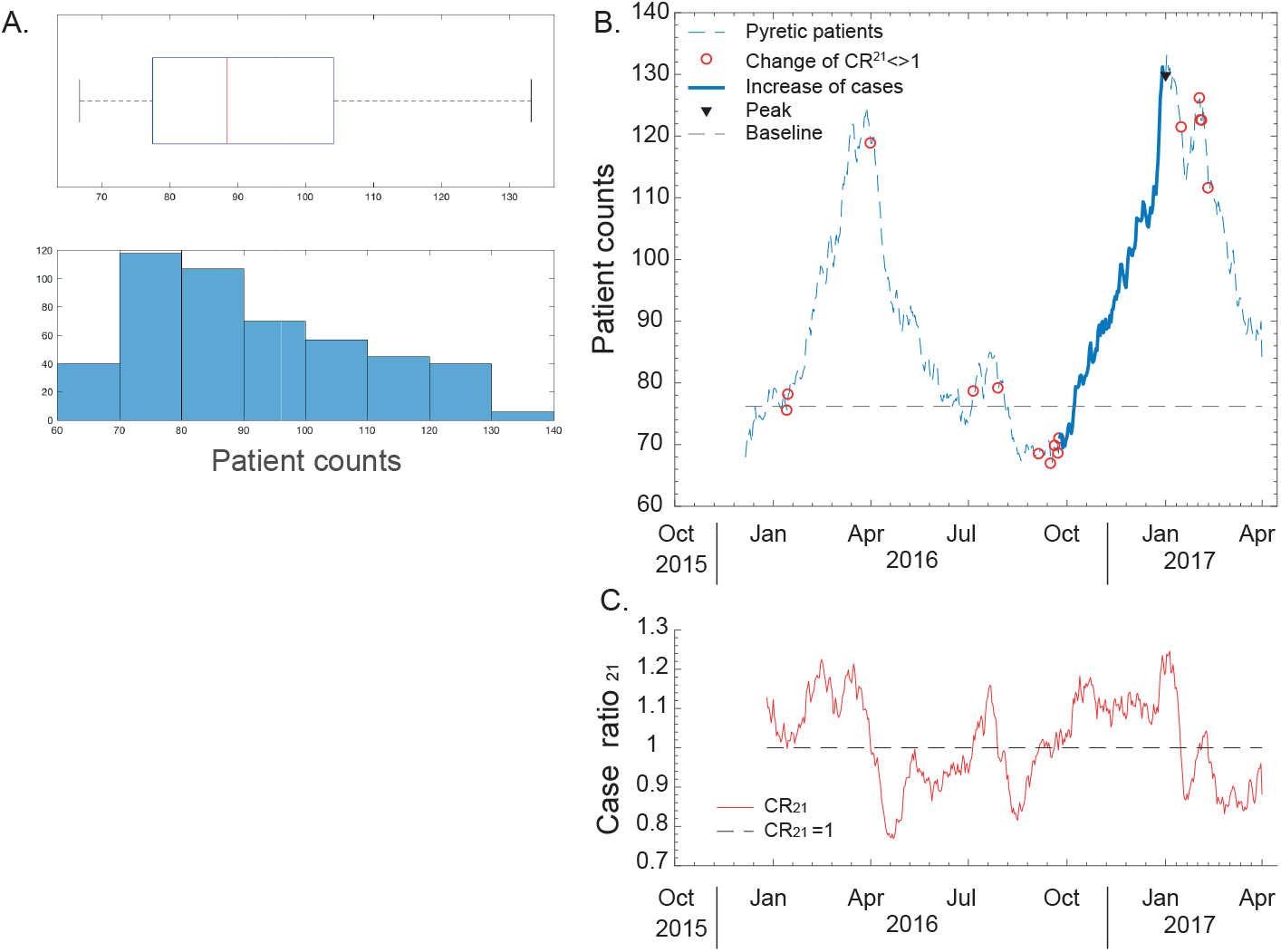
A. histogram and boxplot of daily pyrexia sums. B. daily pyrexia cases. Case Ratio 21 (CR21) indicated as circles. C. CR21 values on the same timescale as B.

To establish the performance of the method, the outbreak detected by PHE was considered the ground truth [31] which allowed to estimate a specificity of 99.7% and sensitivity of 100%. Because of the limited data available the sensitivity and specificity can only be considered crude estimates and should be interpreted accordingly. These can be robustly measure in the future once more specific data is available.

### 3.3 Weekly detection

Temperatures recorded were in range of 21-47° C, with a mean temperature of 36.89° C. As this temperature range is unphysiological it was concluded that some data points were caused by input errors. The temperature-based exclusion removed 8 patients (0.0023%) with temperatures of >42° C, 3 of those within the 42-42.2° C range. The lower temperature cut-off removed 98 patients (0.028%) with temperatures <32° C.

To establish whether seasonal influenza was detectable in the dataset, weekly case numbers were compared with weekly sentinel influenza cases recorded by the ECDC in England (see Figure 3B). Sentinel surveillance data is based on a network of selected health care facilities which select patients with symptoms suggesting influenza for laboratory confirmation. The non-sentinel surveillance is a passive system using patient samples for laboratory confirmation of a variety of sources which are not necessarily from patients showing symptoms of an influenza infection [30].

Both peaks seen for the influenza season 2015/16 and 2016/17 correspond to the data collected by the ECDC. The comparison to the non-sentinel data shows an earlier peak of the ePCR data in the 2016/17 flu season.

This could be explained by the nature of the non-sentinel data, which includes routine patient samples, because the screening is non-targeted.

The ePCR data is mirrored in both the sentinel and non-sentinel laboratory confirmed influenza cases. The ePCR data has a stronger similarity to the non-sentinel surveillance systems which could be explained because all patients are included regardless of symptoms which is also true for the non-sentinel surveillance system.

The smoothed daily pyretic case numbers peaked on the 02/01/17 (week 1/2017). The peak was reached with 133 (18.47%) patients of 721 calls (fractions are caused by the smoothing process using the EMA).

The start of the increase in pyrexia cases defined by a final change of the CR_21_ to >1, was detected on the 24/09/16 (week 38/2016) with 70 pyretic patients (11.2% of 633 calls, Figure 4). This value is 6.8% below the baseline (76.2) and within the standard deviation (23.48), which would not be detectable using a threshold method. This start of the seasonal increase of infections was detected earlier than influenza cases by the ECDC, which identified the start in week 46/2016 [32]. However, this might be due to the fact that the sentinel surveillance runs between October and March and thus had not started when our system detected an increase in cases [33]

To establish the performance of the method, the outbreak detected by PHE was considered the ground truth [31] which allowed to estimate a specificity of 99.7% and sensitivity of 100%. Because of the limited data available the sensitivity and specificity can only be considered crude estimates and should be interpreted accordingly. These can be robustly measured in the future, once more specific data is available.

The weekly case numbers peaked in week 51/2016 with 1042 (21.64%) pyretic patients of 4814 total calls. An increase of the CR_21_ to >1 was detected in week 39/2016 with 570 (13.73%) cases of 4151 calls (Figure 5). The pyrexia count is 5.26% below the baseline (550) at the time of detection and within the standard deviation (134.56), hence would be difficult to detect with a threshold-based method.

**Figure 5:**
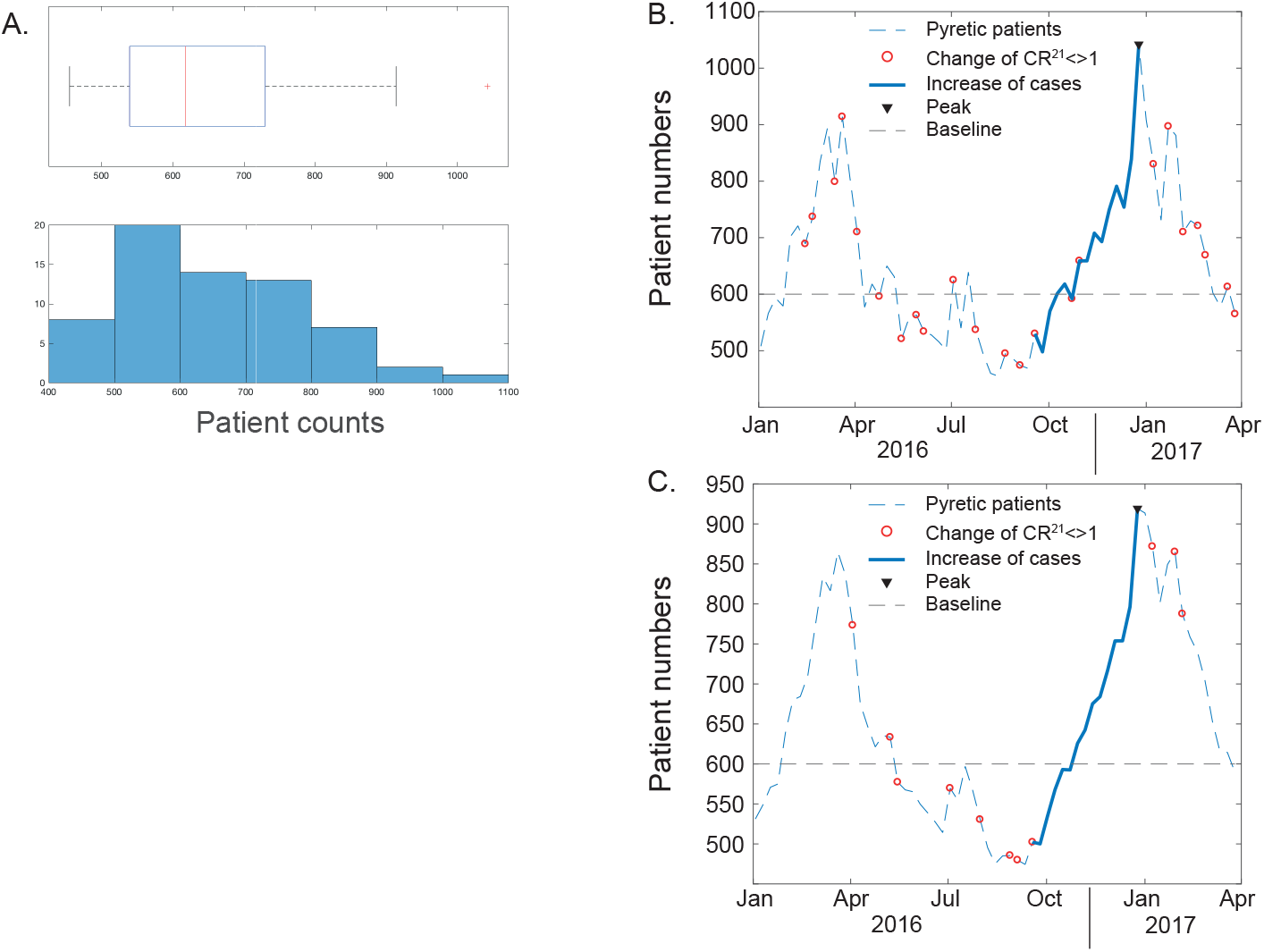
A. histogram and boxplot of weekly pyrexia sums. B. pyrexia unsmoothed cases. Case Ratio 21 (CR21) indicated as circles. C. weekly smoothed (3 week exponential weighted moving average) pyrexia cases. CR21 indicated as circles.

As comparison, the ECDC recorded the first cases in week 46/2016 [32] corresponding to 693 (15.7%) cases of 4407 patients attended by ambulances. This pyrexia count at the date of detection by the ECDC is 15.5% above the baseline (550) but again within the standard deviation (134.56).

The weekly detection method performed with a specificity of 98.2% and sensitivity of 100% when applied to weekly sums and based on the outbreaks as defined by PHE. Again as described above this is based on limited data and only an estimated performance.

### 3.4 Improving accuracy

The weekly data was smoothed using the EMA of 21 days (or 3 sample points), before the sliding CR_21_ was calculated (Figure 5). The smoothed weekly dataset peaked in week 51/2016 with 919 (21.64%) pyretic patients of 4814 total calls. The final CR_21_ change to >1 indicating the start of the outbreak was reached in week 44/2017 with 485.5 (12.5%) pyretic patients of 3868 calls. This case number is 18.40% below the baseline (565) and within the standard deviation (122.33). Once again, the outbreak could not be detected using a threshold method as the number is below the baseline (565) as well as the mean (644.87). Smoothing of the weekly data slightly improved specificity to 99.1% with sensitivity remaining at 100%.

### 3.5 Identification of unknown outbreak during summer 2016

A peak not matching the reference dataset was noticed starting in July 2016 and disappearing by August 2016. It was detected using the weekly method in week 26/2016 and in week 27/2016 by the daily summed method. The CR_21_ recovered to <1 by week 29 and 30/2016 for the weekly and daily method respectively.

This event had small numbers of patients and a short duration not matching the ECDC influenza data. However, this peak correlates with reports from PHE of an *E. coli* outbreak caused by contaminated salad (Figure 6) [34]. The peak in the ePCR data slightly precedes the peak of *E. coli* bactaeremia recorded by PHE [35]. This can be explained by the difference in sampling rate because the data provided by PHE is reported monthly whereas our data is collected daily. Furthermore, the PHE data covers England whereas our data is limited to Devon and Cornwall, which could account for a different timescale of the progression of the outbreak. The outbreak is detected with a delay compared to the PHE data, presumably due to its rapid progress and small patient numbers. A reduction of the delay could be reduced by modifying CR_21_ and applying a smaller *d*.

**Figure 6:**
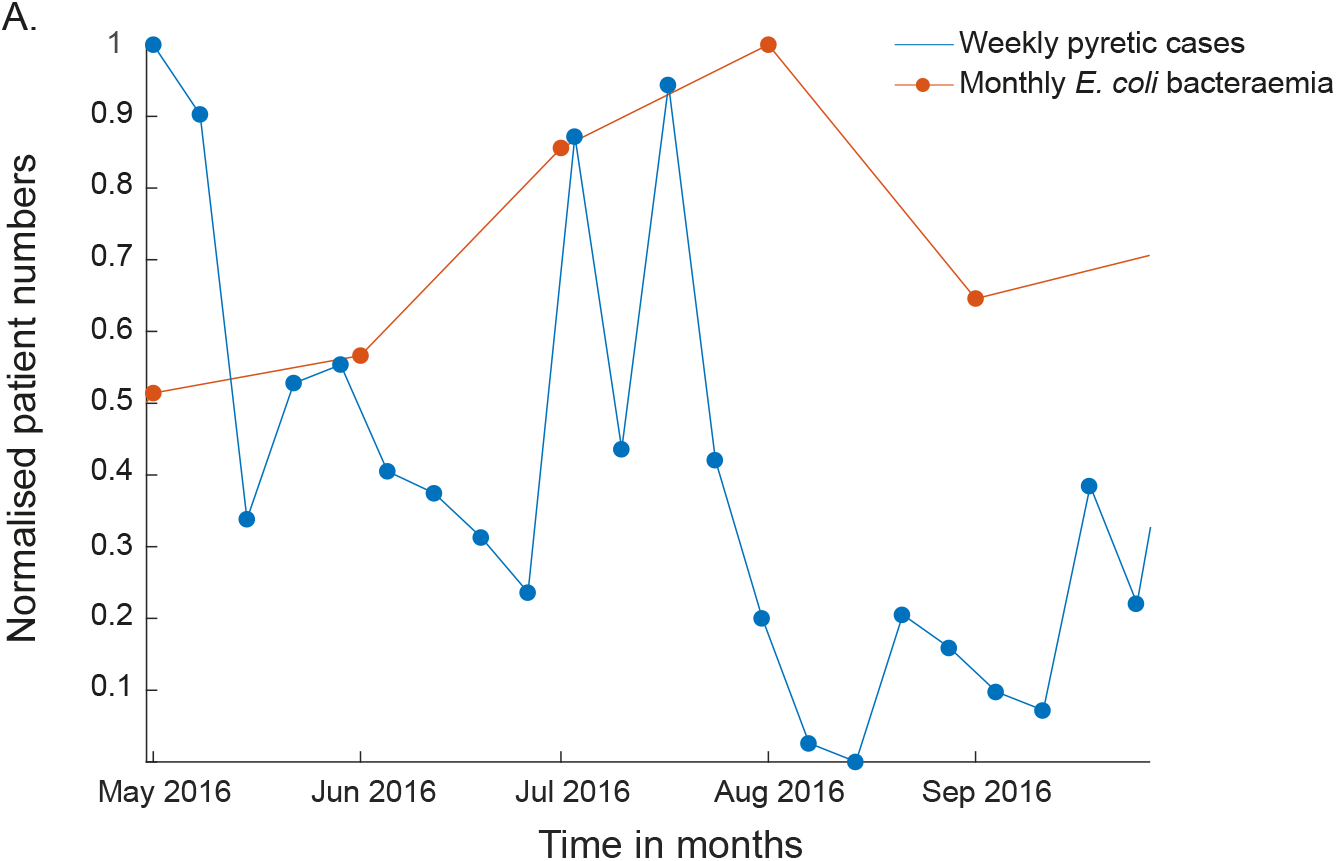
Comparison of weekly pyrexia cases with monthly E. coli bacteremia during the unidentified peak in summer 2016.

The detection of the event caused by *E.coli* shows that the CR_d_ method allows to recognise, not only expected outbreaks, but also events caused by unknown infections.

The outbreak was attributed to the *E. coli* strain O157 and according to PHE reached a final case number of 164 cases with a specific focus in the South-West of England [34,36]. Although the data shown is not specific to strain O157 but shows *E. coli* in general, it suggests that there were considerable numbers of unidentified associated infections. An explanation for this discrepancy could be that only patients with severe symptoms were tested for *E. coli* whereas patients with mild symptoms might have contacted the non-emergency line 111 and were then referred to the ambulance service.

## 4 Discussion

Ambulance crews within SWASFT have collected data for 15.96% of the population in Devon and Cornwall within a year [29]. It must be noted that this includes patients with possible multiple ambulance attendances a year, which is something that cannot be accounted for, as no patient identifying data had been extracted.

The collected data mirrored the gender characteristics of the population although with a focus on the elderly and infants. From these data, it was possible to establish that the pyrexia counts timely matched the seasonal influenza outbreak recorded by the ECDC. As discussed it is reasonable to assume that the majority of the pyrexia cases were due to seasonal influenza but some proportion will have been caused by other circulating infections. This will be addressed in the future by applying the method to more specific syndromes.

Generally epidemiological data tend to be Poisson distributed. However, our data was not Poisson distributed (P values), this could be explained by the fact that only 1.5 seasons were available and would have to be addressed in future studies [9].

Furthermore, using the same data an additional peak was detected, linked to an *E. coli* outbreak concentrated in the South-West of England. The presented method using a sliding CR_d_ reliably detected disease outbreak solely based on tympanic temperature readings. Its non-specificity allowed to detect seasonal influenza and *E. coli* bacteraemia using the same data. This demonstrates the ability of this method to detect outbreaks of unknown causes.

The seasonal increase of fever cases was detected up to 9 weeks before influenza cases were recorded by conventional methods employed by the ECDC. As the method described here does not specifically detect influenza several contributing factors can be attributed to this finding. In the UK the sentinel detection runs between October and March thus it could not detect earlier cases. Furthermore, during winter many different infectious agents are circulating including the common cold, all of which cause fever.

Due to the nature of the described method to date the infectious agent cannot be narrowed down, however this will be addressed by applying it to more specific syndromes.

The demonstrated method performed with a very similar sensitivity and specificity to the method demonstrated by Singh *et al.* 2010 [10]. However, it has to be emphasised that these are estimates and are based on limited data therefore have to be interpreted with caution. Because it does not rely on thresholds set by an investigator, it is ideally suited as an EED system for unknown outbreak causes.

Due to its simplicity, the CR_d_ method is easy to deploy and will effectively detect a wide range of syndromes. By using case numbers of different symptoms, this method will also allow to automatically distinguish between different infections. For example, patients with a combination of pyrexia and diarrhoea could be monitored separately from patients experiencing pyrexia only.

The current shortcoming of the method is that it was only applied to pyrexia data but no other syndromes. It is to be expected that the combination of pyrexia with diarrhoea would have allowed to immediately distinguish between the peak potentially caused by seasonal influenza and the event caused by *E. coli.* Therefore, as a continuation of this study, different symptoms will be combined to allow the monitoring of several syndromes in parallel. Although the proposed method does not rely on a threshold to detect an increase in cases, it still requires the user to define the value of *d*, which will normally require some knowledge about the transmission rate of the monitored infection.

The necessity to choose a time frame raises similar questions as threshold-based methods, on how to decide on the best time frame. Therefore, this flexibility makes the CR_d_ method able to adapt to the data and transmission rates allowing to reduce false positives in noisy data.

In practice this method could be further developed to allow the surveillance of several different syndromes but can also be used to anticipate spikes in ambulance call rates as well as hospital admissions. It might also lead the way to predict the length and magnitude of a detected outbreak.

## 5 Conclusion

It was demonstrated that electronic patient records documented by ambulance crews correlate with the sentinel data collected by the ECDC and that it allows the implementation of an EED system. The CR-method was found to be sensitive to several different infectious agents based on the same data.

The detection of events occurred earlier compared to the ECDC but to date does not allow to distinguish between infectious agents. Therefore, the move to digital patient records allows to monitor the entire population for several syndromes at high sample rates and for several syndromes at the same time, making it an ideal data source for an EED system.

## Footnotes

1 All reference datasets provided by ECDC are based in turn on data provided by WHO and Ministries of Health from the affected countries. The views and opinions of the authors expressed herein do not necessarily state or reflect those of the ECDC. The accuracy of the authors’ statistical analysis and the findings they report are not the responsibility of ECDC. ECDC is not responsible for conclusions or opinions drawn from the data provided. ECDC is not responsible for the correctness of the data and for data management, data merging and data collation after provision of the data. ECDC shall not be held liable for improper or incorrect use of the data.

